# Mapping protein-metabolite interactions in *E. coli* by integrating chromatographic techniques and co-fractionation mass spectrometry

**DOI:** 10.1101/2024.02.14.580258

**Authors:** Mateusz Wagner, Jieun Kang, Catherine Mercado, Venkatesh P. Thirumalaikumar, Michal Gorka, Hanne Zillmer, Jingzhe Guo, Romina I. Minen, Caroline F. Plecki, Katayoon Dehesh, Frank C. Schroeder, Dirk Walther, Aleksandra Skirycz

**Author notes:** shared authorship (Mateusz Wagner, Jieun Kang).

## Abstract

In our pursuit of understanding the protein-metabolite interactome, we introduced PROMIS, a co-fractionation mass spectrometry (CF-MS) technique focusing on biosynthetic and regulatory processes. However, the challenge lies in distinguishing true interactors from coincidental co-elution when a metabolite co-fractionates with numerous proteins. To address this, we integrated two chromatographic techniques— size exclusion and ion exchange—to enhance the mapping of protein-metabolite interactions (PMIs) in *Escherichia coli*. This integration aims to refine the PMI network by considering size and charge characteristics, resulting in 994 interactions involving 51 metabolites and 465 proteins. The PMI network is enriched for known and predicted interactions validating our approach’s efficacy. Furthermore, the analysis of protein targets for different metabolites revealed novel functional insights, such as the connection between proteinogenic dipeptides and fatty acid biosynthesis. Notably, we uncovered an inhibitory interaction between the riboflavin degradation product lumichrome and orotate phosphoribosyltransferase (PyrE), a key enzyme in *de novo* pyrimidine synthesis. Lumichrome supplementation mimicked the biofilm formation inhibition observed in a *ΔpyrE* mutant strain, suggesting lumichrome role in integrating pyrimidine and riboflavin metabolism with quorum sensing and biofilm formation. In summary, our integrated chromatographic approach significantly advances PMI mapping, offering novel insights into functional associations and potential regulatory mechanisms in *E. coli*.

## Introduction

Protein-metabolite interaction (PMI) networks remain largely uncharacterized even in simple model systems, although metabolites play essential role in regulating protein activities, protein-protein interactions, subcellular localization, and protein condensed states (Venegas-Molina *et al*, 2021; Diether & Sauer, 2017; Kosmacz *et al*, 2020; Baker & Rutter, 2023). Small molecule regulation is known for almost all protein classes. Notably, nearly 75% of the 300 transcription factors (TFs) in *Escherichia coli* have predicted small-molecule binding domains (Ledezma Tejeida *et al*, 2021). PMI networks are often complex and highly dynamic, responding rapidly to environmental and developmental cues. Recent studies highlight the potential for a single protein to engage with multiple small molecule ligands under different conditions, facilitating the discovery of previously unknown regulatory connections (Piazza *et al*, 2018; Tian *et al*, 2012; Lim *et al*, 2018; Diether *et al*, 2019; Link *et al*, 2013; Hackett *et al*, 2016; Lempp *et al*, 2019).

Biochemical methods for the identification of PMIs range from the classical affinity purifications that exploit either a metabolite or a protein bait to identify partners to more recent approaches that measure ligand-induced changes in the protein properties such as thermal stability or accessibility to a protease around the binding site (Wagner *et al*, 2021; Luzarowski & Skirycz, 2019; Diether & Sauer, 2017; Venegas-Molina *et al*, 2021; Orsak *et al*, 2012). The methods mentioned above have their unique advantages and drawbacks but depend on availability of a metabolite or protein bait. We previously developed PROMIS <u>(PRO</u>tein-<u>M</u>etabolite Interactions using <u>S</u>ize separation) (Veyel *et al*, 2018), a co-fractionation mass spectrometry (CF-MS) approach for proteome and metabolome-wide mapping of protein-metabolite complexes to circumvent this limitation. CF-MS methods combine separation of native complexes with mass spectrometry-based analysis of the obtained fractions and use the similarity of elution profiles to delineate interactors (Schlossarek *et al*, 2023). CF-MS approaches have been instrumental in generating comprehensive protein-protein interaction networks across model and non-model organisms, e.g., (Skinnider *et al*, 2021; Wan *et al*, 2015; Mallam *et al*, 2019; Havugimana *et al*, 2012; Lee & Szymanski, 2021; Lee *et al*, 2021; Aryal *et al*, 2014; McWhite *et al*, 2020). In its original implementation, PROMIS relied on size separation chromatography (Veyel *et al*, 2017); metabolites bound to protein complexes thus would separate into earlier-eluting high molecular weight fractions, whereas unbound small molecules separate into late-eluting low molecular weight fractions. Fractions are then subjected to proteomic and metabolomic analysis, and the resulting metabolite and protein profiles are searched for co-eluting metabolites and proteins. Recent PROMIS studies in Arabidopsis (Veyel *et al*, 2018) and yeast (Schlossarek *et al*, 2022; Luzarowski *et al*, 2021) reported hundreds of metabolic features separating with protein complexes, of which only a subset could be annotated and many more awaiting chemical identification. Independently and in an analogous SEC-based CF-MS experiment, Li and colleagues (Li *et al*, 2021) reported 461 putative interactions in the thermophilic fungus *Chaetomium thermophilum*, including a novel regulatory pairing between isopentenyl adenine and the ribosome.

However, in CF-MS experiments a single metabolite often co-fractionates with hundreds of different proteins, of which only a subset may represent “true interactions”, while the presence of most other proteins is due to coincidental co-elution. Identifying the true binders represents the most significant challenge in using CF-MS-based methods for mapping PMI networks. Based on previous work on protein-protein interactions (Schlossarek *et al*, 2023), we hypothesized that integrating PROMIS with two orthogonal chromatographic methods may greatly improve the ability to distinguish between true and coincidental co-elution. As a second chromatography to complement SEC we specifically focused on ion exchange (IEX), given that many similarly sized protein complexes will differ in charge and *vice versa*. Separation of protein drug complexes has been previously demonstrated by Chan and colleagues. Their method, termed TICC (<u>T</u>arget Identification by <u>C</u>hromatographic <u>C</u>o-elution), relies on the distinctive shift observed in a small molecule’s chromatographic elution profile upon binding to its protein target (Chan *et al*, 2012). Chan and colleague’s experimental strategy encompassed two native IEX separations of the cell lysate incubated with the studied compounds and a control sample containing just the drug that enabled differentiating between the elution profiles of free drug versus drug-protein complex. More recently, TICC was used to separate antibiotic-protein complexes from bacteria (Schäkermann *et al*, 2021).

The central objective of our research presented in this study is to assess whether integrating IEX (TICC) and SEC (PROMIS) can enhance the accuracy of mapping endogenous PMI networks in cell lysates. For this purpose, we selected *E. coli* due to the relatively low complexity of its proteome and metabolome, coupled with the abundance of known or predicted interactions available from databases such as STITCH (Kuhn *et al*, 2010). As anticipated, we found that integration of PROMIS and TICC facilitates distinguishing between true and coincidental co-elution events, greatly improving our ability to map PMI networks.

## Results and Discussion

### Ion exchange is suitable for the separation of protein-metabolite complexes from complex cell lysates

This study aimed to test whether using two orthogonal chromatographies, size exclusion (SEC), and ion exchange (IEX), could significantly increase the scope of CF-MS for delineating PMI networks. Our overall experimental strategy is outlined in **Figure 1a-c**. Analogous to previous studies (Veyel *et al*, 2018; Luzarowski *et al*, 2021), PROMIS separation using SEC of *E. coli* lysate yielded 40 protein-containing fractions spanning proteins and protein complexes from approximately 10 MDa to 10 kDa. The collected SEC fractions were analyzed by untargeted liquid chromatography-mass spectrometry (LC-MS) metabolomics and LC-MS/MS proteomics. In a separate experiment, we subjected the same lysate sample to IEX fractionation (IEX_S fraction set). To distinguish between metabolite peaks corresponding to free *vs.* protein-bound metabolites, we additionally applied the same IEX fractionation protocol to heat-denatured lysate (IEX_C fraction set). The IEX_S separation yielded 24 fractions of which 17 contained proteins, which were subjected to untargeted metabolomics and proteomics. SEC and IEX separations were performed for two independent experiments yielding four CF-MS datasets. Metabolic features were annotated by using a library of authentic standards. The comparison of metabolite elution profiles between IEX_C and IEX_S focused on eliminating peaks representing unbound metabolites, as determined by the control separation (**Figure 1d-e**). Note that, although protein-bound metabolites were identified primarily based on shifts of their elution maxima in IEX_S *vs.* IEX_C, we also considered metabolite peaks whose intensity was strongly increased compared to the control separation as potentially protein-bound (**Figure 1e**, see example of niacinamide). The profile of differential metabolite elution comparing IEX_S and IEX_C we refer to as the omniTICC profile (Chan *et al*, 2012). The “omni” points to the untargeted nature of the approach, enabling characterization of endogenous protein-metabolite complexes.

**Fig 1.**
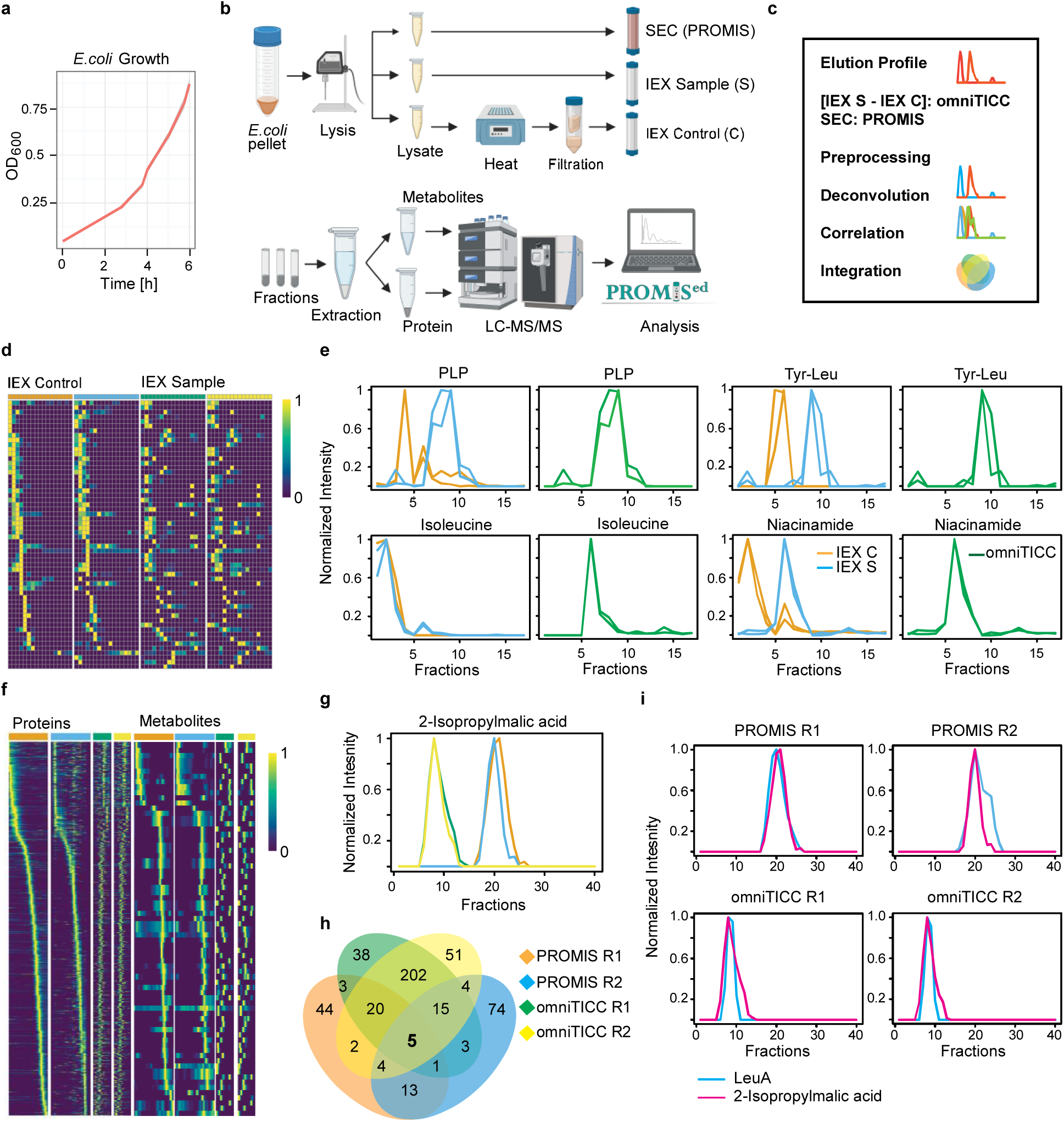
Overview of the experimental strategy. a) *E. coli* growth curve. Bacterial cells were collected at the exponential phase (OD_600 nm_ = 0.8). b, c) Schematic representation of the experimental strategy and data analysis steps. d) Heat-map representation of the elution profile of the metabolites across examined fractions in the IEX control and IEX sample separations. e) Example elution profiles of four representative metabolites obtained from IEXs and omniTICC. f) Heat-map representation of the protein and metabolite elution profiles across PROMIS and omniTICC separations from two independent experiments. g) A representative example of an elution profile of a metabolite, 2-isopropylmalic acid, in the PROMIS and omniTICC separations from two independent experiments. h) Proteins co-eluting with 2-isopropylmalic acid in the four separations (Pearson correlation coefficient > 0.8). i) Co-elution of 2-isopropylmalic acid and its well-known interactor, LeuA, d-i) R1 and R2 stand for two independent biological replicas.

Proteomics and metabolomics of the SEC and IEC fractions revealed 1479 proteins and 58 known metabolites, annotated using a compound reference library (**Figure 1f; Supplementary Dataset S1-S4**). Functional analysis of the 1479 proteins revealed enrichment of categories broadly associated with metabolism. The 58 annotated metabolites included amino acids, nucleotides, cofactors, dipeptides, and cyclic dipeptides as well as diverse metabolic intermediates. Our results demonstrate that IEX can be effective at separating protein-metabolite complexes within a complex cellular lysate, complementing SEC.

### Functional insight from global analysis of the protein-metabolite interactome

To construct the protein-metabolite interactome of *E. coli*, we looked for protein-metabolite pairings that co-elute in both the PROMIS and omniTICC separations, across two independent repeats. Pre-processing, deconvolution, correlation, and integration steps were performed using the standard analysis settings embedded in the PROMIsed app (Schlossarek *et al*, 2021). An example analysis is outlined in **Figure 1g-I** for isopropylmalate, an intermediate in leucine biosynthesis. In a single PROMIS or omniTICC separation, isopropylmalate co-eluate ((Pearson correlation coefficient (PCC) >0.8)) with tens of proteins, but only five proteins co-eluate (PCC >0.8) across all four datasets (two repeats of SEC and IEC, **Figure 1h**). Among these five putative protein targets, LeuA, an enzyme involved in leucine synthesis, is a known isopropylmalate binder (**Figure 1i**).

Analogous analysis was performed for all 58 metabolites to generate a global map of the *E. coli* PMI network. The resulting network encompasses 51 metabolites, 465 protein nodes, and 994 PMIs (**Figure 2a; Supplementary Dataset S5**). The 0.8 PCC cut-off to delineate putative interactors was determined by optimizing the true positive (sensitivity, true targets) to false positive (1 minus specificity, coincidental coelution) ratio. This optimization was based on the evaluation of 1012 known or predicted interactions retrieved from the STITCH database (referred to as STITCH interactions) between the 58 metabolites and 1479 proteins in our dataset. The network recapitulated 92 of the STITCH interactions, which is 7.4 times more than expected by chance, based on the results of the positive likelihood ratio (sensitivity/(1-specificity)) (**Figure 2b**). In comparison, using PROMIS or omniTICC alone captures approximately two times more STITCH interactions than expected by chance (**Figure 2b**). To learn which compounds drive the positive likelihood ratio, nucleotide monophosphates (NMPs), nucleosides, amino acids, and cofactors were examined separately (**Figure 2b**). The PROMIS & omniTICC-derived PMI network outperformed PROMIS or omniTICC alone in recapitulating STITCH interactions for NMPs and amino acids but not for cofactors (**Figure 2b**). Neither the single nor combined approaches captured more nucleoside STITCH-retrieved targets than expected by chance. Close inspection of the data revealed that, unlike amino acids, NMPs, and nucleosides, the elution profiles of cofactors, particularly nicotinamide adenine dinucleotide (NADH) and flavin adenine dinucleotide (FAD), differed between the two PROMIS separations (**Figure S1a**). When we considered proteins cofractionating with cofactors in the two omniTICC separations but only in one of the two PROMIS experiments, we found that the positive likelihood ratio increased from 0.89 to 2.3 (**Figure 2b**); an improved performance compared to PROMIS or ominiTICC alone.

**Figure 2.**
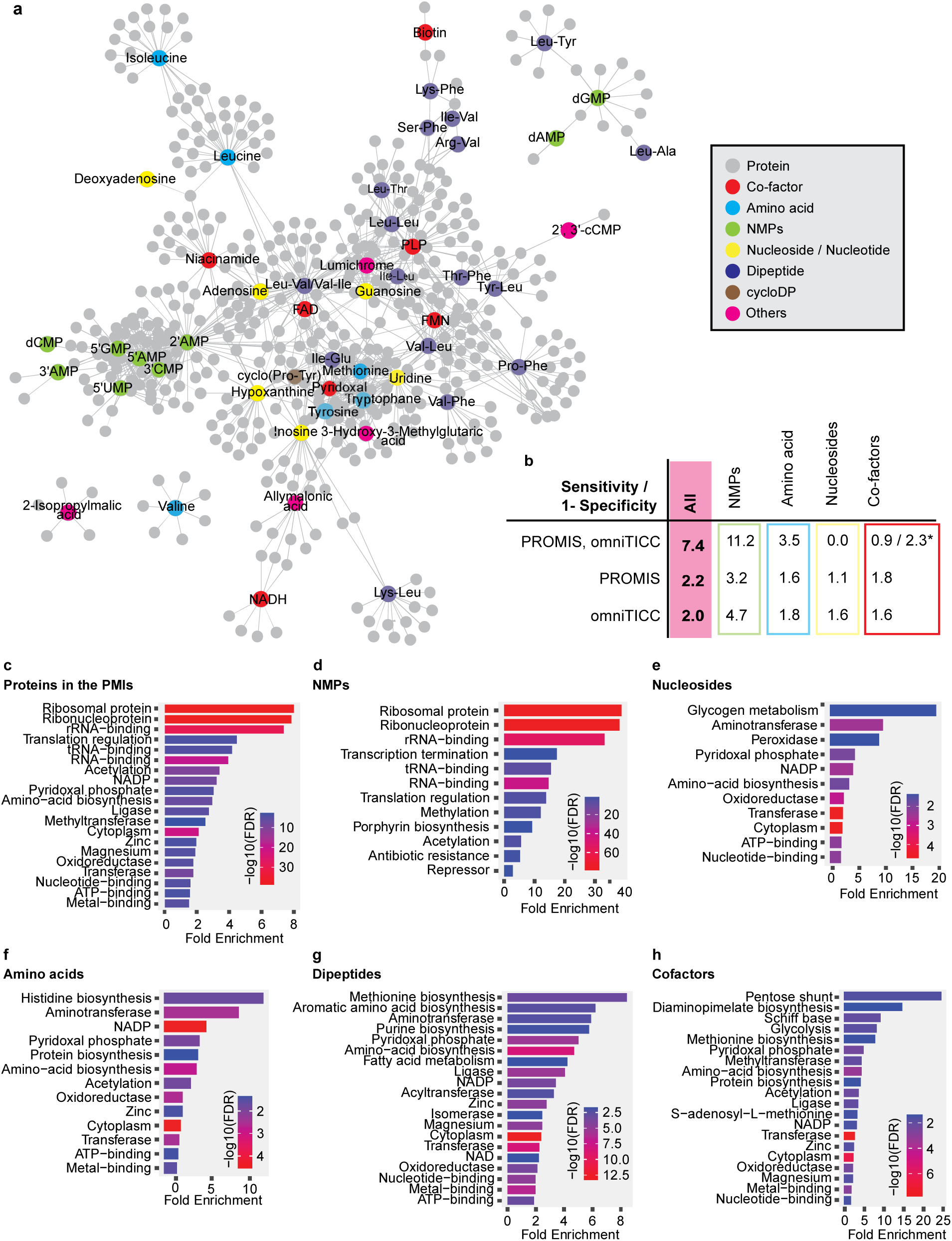
*E. coli* PMI network derived from integration of PROMIS and omniTICC separations. a) The edges between metabolites and proteins derive from co-elution. b) Known PMIs deposited in the STITCH database (confidence score >0.4, based on the experimental evidence) were used to calculate sensitivity/1-specificity ratio for the entire network, and subnetworks of the different compound classes. Co-factors* - edges derived from the two omniTICC, but just a single PROMIS separation. c-h) Functional over-representation of the protein targets of the different compound classes was calculated and visualized using SinyGO 0.77 app (Ge *et al*, 2020).

Functional analysis of the 465 protein nodes captured in the network revealed enrichment of categories broadly related to translation and cellular metabolism, particularly amino acid biosynthesis and carbon metabolism (**Figure 2c**). Specifically, the enrichment of proteins associated with translation was driven by interactions between NMPs and ribosomes (**Figure 2d**). In contrast to NMPs, which clustered together, the five nucleosides: guanosine, adenosine, uridine, inosine, deoxyadenosine, and nucleotide hypoxanthine had distinct sets of putative targets (**Figure 2e**). Among these proteins are several transcription factors, proteins involved in RNA modifications, and enzymes of glycogen metabolism. In addition to acting as metabolic intermediates, nucleosides are known for their regulatory and signaling functions, and hence it is expected that they may have multiple and specific protein targets. For instance, hypoxanthine binding to the transcription factor PurR enhances repression of its target genes, thereby affecting the metabolism and transport of purine and pyrimidine nucleotides (Cho *et al*, 2011). The PuR and hypoxanthine interaction was also captured in PROMIS and omniTICC separations (**Figure S1b**).

Corroborating our previous work in plants and yeast (Veyel *et al*, 2018; Luzarowski *et al*, 2021; Schlossarek *et al*, 2022), the PMI network contained multiple proteinogenic dipeptides. With some exceptions, dipeptides are typically products of protein degradation (Minen *et al*, 2023; Thirumalaikumar *et al*, 2020; Tsukahara *et al*, 2018). Notably, of the 17 dipeptides in the network, 13 have at least one branched chain amino acid, valine, leucine, or isoleucine (**Figure 2a**). The various dipeptides, including the retro dipeptides such as Tyr-Leu and Leu-Tyr, have a unique set of putative targets distinct from those delineated for individual amino acids, including valine, leucine, or isoleucine (**Figure 2a**). Distinct activities of dipeptides and amino acids have been reported in previous studies (Minen *et al*, 2023; Cheng *et al*, 2023; Ichinose *et al*, 2015; Mizushige *et al*, 2020; Zhang *et al*, 2016; Naka *et al*, 2015). Dipeptides were found in complexes with enzymes of central carbon metabolism in plants and yeast (Moreno *et al*, 2021; Luzarowski *et al*, 2021). Our *E. coli* dipeptide-enzyme interaction network is enriched in enzymes of amino acid metabolism, particularly aromatic amino acids, and lysine biosynthesis (**Figure 2g**). Given that dipeptides act as a read-out of nitrogen status and a source for amino acids, their association with amino acid synthesis is unsurprising (Tegeder & Rentsch, 2010) . Furthermore, the concept of end-product inhibition in amino acid synthesis, well-described in bacteria for regulating amino acid levels (Chubukov *et al*, 2014), leads us to speculate that dipeptides may serve analogous functions. Additionally, dipeptide targets include dipeptide transporter (DppA) and oligopeptide transporters (OppA and SapA). Dipeptide regulation of peptide transport has been reported before in yeast (Xia *et al*, 2008; Du *et al*, 2002). Finally, the lists of putative targets for amino acids include expected categories such as amino acid biosynthesis enzymes (**Figure 2f**), whereas the molecular terms overrepresented among putative targets of co-factors include “pyridoxal phosphate” and “NADP binding” and encompass multiple enzymes of carbon and amino acid metabolism (**Figure 2h**).

These results demonstrate that integrating two chromatographic techniques in a co-fractionation mass spectrometry approach can enhance PMI mapping capabilities and that resulting PMI networks may provide broad functional insights.

### Identification of novel regulatory PMIs in lipid and nucleotide metabolism

The *E. coli* PROMIS and omniTICC-derived interaction network presented here contains nearly a thousand putative interactions. One way to prioritize PMIs for binding and functional characterization is to select associations that may have escaped traditional binding screens usually biased toward well-established small molecule effectors. Dipeptides are good examples of compounds reported to have diverse, only recently uncovered bioactivities; however, they remain understudied (Minen *et al*, 2023). Protein interactors of dipeptides in our network comprised multiple enzymes, also including enzymes associated with fatty acid biosynthesis. This observation may suggest a regulatory interplay between protein degradation and lipid metabolism. To follow up on this intriguing hypothesis, we selected a representative interaction between a dipeptide, Val-Leu, and FabF (**Figure 3a**), which catalyzes the conversion of palmitoyl-ACP to 3-keto-cis-vaccenoyl-ACP, a crucial step in the type II fatty acid elongation cycle (Edwards *et al*, 1997). To investigate potential binding between FabF and Val-Leu, we employed microscale thermophoresis (MST). MST exploits the differential movement of the free ligand versus ligand-protein complex in a microscopic temperature gradient (Jerabek-Willemsen *et al*, 2011). The measured binding constant (K_d_) of 12-14 μM between FabF and Val-Leu (**Figure 3b-c**) falls within the range of reported dipeptide concentrations (Heidenreich *et al*, 2021). Only one binding pocket was predicted with high confidence for FabF, using computational binding site prediction (see Methods), coinciding with the binding site for the co-crystallized ligand platencin in the PDB structural record for FabF (PDB-ID 3HO9) (**Figure 3d-f**). We used this binding pocket for molecular docking analysis and predicted Val-Leu binding with binding affinity within the range of nM to µM, in reasonable agreement with the experimentally determined binding constant.

**Figure 3.**
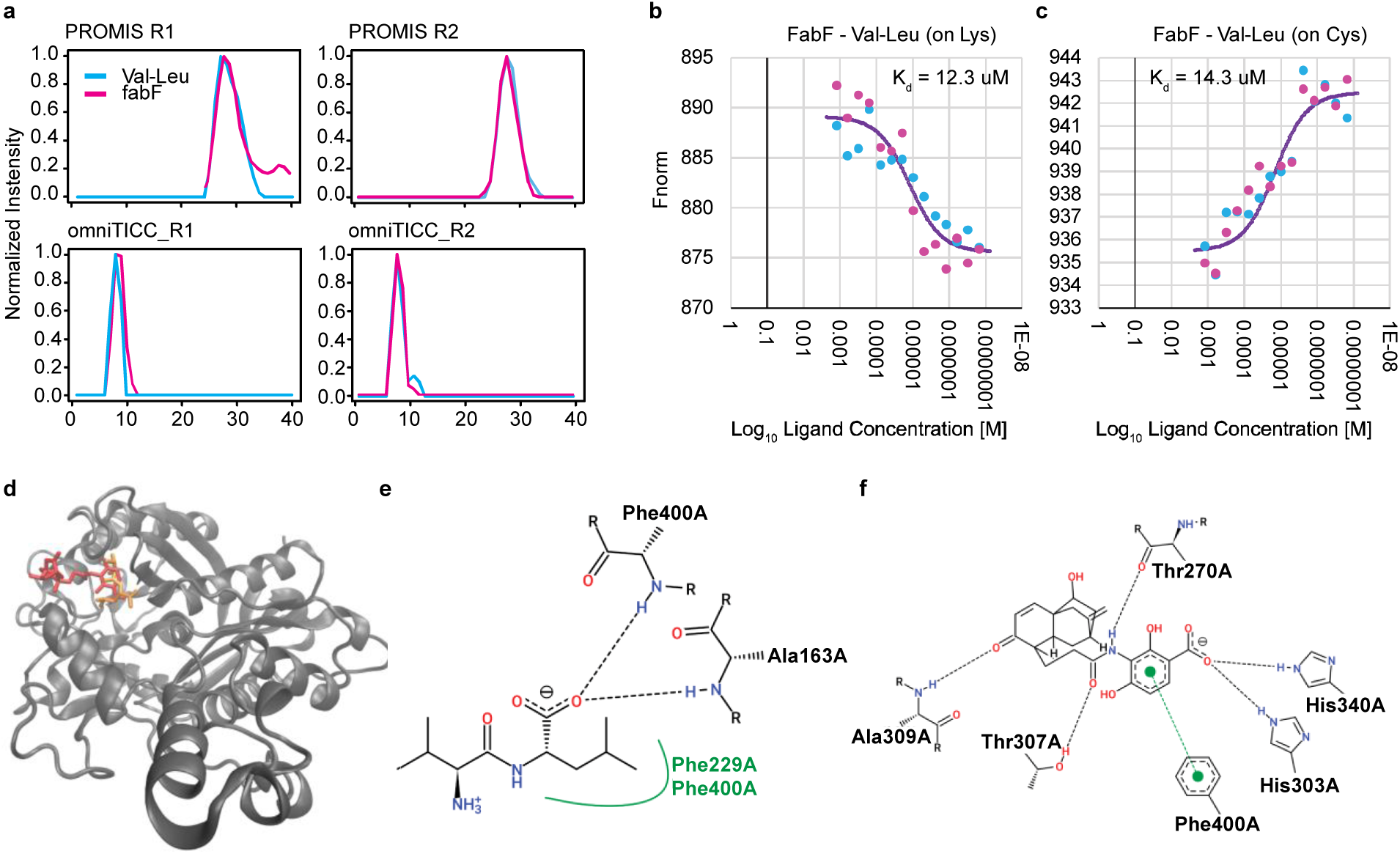
Novel binding events derived from the PMI network. a) Co-elution profiles between dipeptide Val-Leu and enzyme FabF. b, c) MST binding curves; ligand corresponds to Val-Leu. The FabF protein was labeled using two different dyes, NHS dye that labels lysine and NHS-Maleimide dye that labels cysteine. Each binding curve is derived from two independent titrations and was fitted using Monolith analysis software. d-f) Potential interaction mode between FabF (PDB ID: 3HO9) and Val-Leu (PubChem ID: 6993118) obtained via molecular docking. d) Ribbon diagram of the 3D-structure of FabF with the reference (co-crystallized in PDB file) ligand, platencin (PubChem ID: 16745128), in red, and the top docking pose of Val-Leu in orange. The predicted binding affinity for this top docking pose was estimated to lie in the nM to µM range, and within a nM range for platencin. e-f) 2D ligand-protein interaction diagrams of FabF and Val-Leu e) and platencin f). The dashed lines represent hydrogen bonds, while the green spline segment represents hydrophobic contacts between Val-Leu and FabF.

Another strategy that can be employed to prioritize interactions builds on the known conservation of regulatory PMI (Gruber *et al*, 2021). Hence, we wondered if we could tap into previously obtained PROMIS datasets from plants and yeast to assist in prioritizing PMIs from *E. coli* based on presumed conservation. As a proof-of-concept example, we chose the photoconversion product of the cofactor riboflavin, lumichrome (Treadwell & Metzler, 1972), which we noted was present in the PROMIS separations from *E. coli*, yeast (Luzarowski *et al*, 2021) and Arabidopsis (Veyel *et al*, 2018). Using a less stringent PCC cut-off of 0.7 to define coelution, we found that five of the 82 putative lumichrome targets in *E. coli* had a homolog in yeast and Arabidopsis that was also co-fractionating with lumichrome (**Figure 4a**). Of those five proteins, two had been previously associated with a related flavin molecule, the co-factor flavin mononucleotide (FMN) (**Figure 4b**), and we decided to focus on one of them, the enzyme orotate phosphoribosyltransferase (PyrE) (**Figure 4c**). Notably, the riboflavin and *de novo* pyrimidine synthesis pathways are biochemically connected, as they both rely on the intermediates of the pentose phosphate pathway. PyrE plays a pivotal role in *de novo* uridine monophosphate (UMP) biosynthesis by catalyzing a reaction between orotic acid and 5-phosphoribosyl-1-pyrophosphate (PRPP), ultimately yielding orotidine monophosphate (OMP) (Kim *et al*, 2003). In eukaryotes, the homolog of pyrE is called uridine monophosphate synthase (UMPS) and combines the activities of two enzymes, orotate phosphoribosyl transferase and orotidine-5’-decarboxylase. To test if lumichrome interacts with purified PyrE protein, we again employed MST. A control binding assay confirmed the interaction between PyrE and its well-known ligand PRPP, with a K_d_ value of 8.21 μM (**Figure 4d**), and we demonstrated binding between PyrE and lumichrome, with a binding affinity of 90.3 μM (**Figure 4d**). To further confirm PyrE interaction with lumichrome, we conducted a thermal shift assay (TSA). This method relies on the premise that ligand binding often influences the melting temperature of its protein target. We measured a small but reproducible shift in the melting temperature of PyrE according to an increase in the concentration of the ligands within the range of 74.17 – 74.76 °C for PRPP and 74.17-74.91 °C for lumichrome (**Figure 4e**). These observations support that lumichrome in fact binds to PyrE.

**Figure 4.**
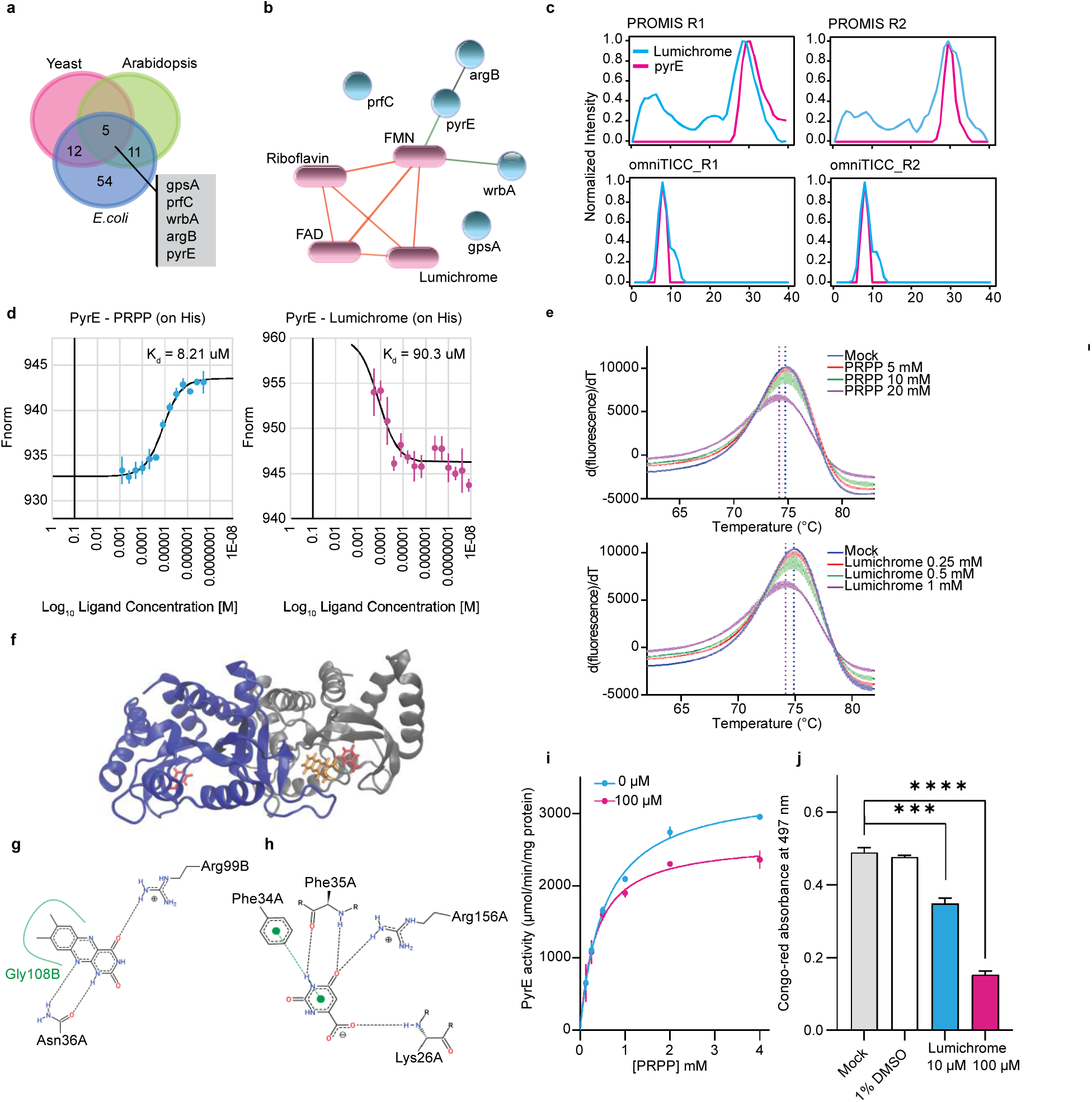
Novel binding events derived from the PMI network. a) Venn representation focused on the overlap of *E. coli* putative lumichrome targets with proteins co-eluting with lumichrome in *A. thaliana* (Veyel *et al*, 2018) and *S. cerevisiae* (Luzarowski *et al*, 2021) in PROMIS experiments. Homology was derived using DIOPT ortholog prediction tool (Hu *et al*, 2011). b) The STITCH derived interactions between the five proteins co-eluting with lumichrome in bacteria, yeast and plants with the different flavin compounds. Edges are based on the experimental, database and literature evidence with the confidence score >0.4.c) Co-elution profiles between lumichrome and enzyme PyrE. d) MST binding curves for PyrE-PRPP and PyE-lumichrome. PyrE was labeled at the position of His-tag. Each curve was generated by three independent titrations with two replicates. e) Melting curves of PyrE upon treatment with PRPP and lumichrome. Each curve was derived from three replicates. f-h) Potential interaction mode between PyrE (PDB ID: 6TAK) and lumichrome (PubChem ID: 5326566) obtained via molecular docking. f) Ribbon diagram of the 3D-structure of PyrE. Chain A of PyrE is shown in grey and chain B in blue. The reference (co-crystallized in PDB file) ligand, orotate (PubChem ID: 1492348), and the top docking pose of lumichrome are shown in red and orange, respectively. The predicted binding affinity for this top docking pose was estimated to lie in the µM to mM range, and within a nM range for orotate. g-h) 2D ligand-protein interaction diagrams of PyrE and lumichrome (g) and orotate (h). The dashed lines represent hydrogen bonds, while the green spline segment represents hydrophobic contacts between the lumichrome and PyrE. i) PyrE activity upon treatment with 100 μM of lumichrome. The activity was determined by calculating the amount of enzyme required to convert 1 μmol of orotic acid to orotidine monophosphate (OMP) per minute (n=3). j) The inhibitory effect of lumichrome on congo-red binding to biofilm components. Bacterial supernatant was measured at 497 nm after incubating with lumichrome for 24 h (n=3).

Using the PDB entry 6TAK (excluding the co-crystallized ligand orotate) as the target protein structure, *in silico* molecular docking to the pocket with the highest probability predicted lumichrome binding within an affinity range of µM to mM (**Figure 4f-h**). Of note, while the predicted binding site encompasses the binding pocket of orotate, lumichrome was predicted to bind to the larger pocket adjacent to the minor pocket occupied by orotate. Interestingly, orotate and lumichrome share an interaction edge (O=C-NH-C=O, **Figure 4f**), suggesting a similar binding mode. However, lumichrome is larger and does not fit into the orotate pocket. Visual inspection of the 3D structure reveals that a “lid” formed by a short beta-sheet segment may swing open, allowing lumichrome to bind in place of orotate. However, the available apo-structure (6TAI) does not show this lid as open, and enzymatic analysis (see below) argues against a competitive binding. An extended molecular docking simulation with a flexible protein backbone position will be necessary to investigate the binding details further.

After confirming the binding of lumichrome to PyrE, we investigated the impact of this binding on intrinsic PyrE activity. A continuous spectrophotometric method was used to monitor the activity of OPRTase in presence of PRPP and orotate. This method measures decreasing absorbance of orotate at 295 nm as OPRTase catalyzes conversion of orotate into OMP (Scapin *et al*, 1995). Enzyme activity reached saturation as the concentration of PRPP increased from 0.125 mM to 4 mM (**Figure 4i**). The presence of 100 μM of lumichrome reduced PyrE activity by approximately 20 % at a concentration of 4 mM. The minimum tested lumichrome concentration that significantly decreased PyrE activity was 25 μM, whereas an increase in lumichrome concentration above 100 μM and up to 400 μM did not yield a stronger inhibitory effect (data not shown). The inhibitory effect of lumichrome on PyrE prompted us to investigate the effect of lumichrome on biofilm formation, since the *de novo* nucleotide biosynthesis pathway is closely linked to production of curli amyloid fibers, which constitute the extracellular matrix of biofilms in *E. coli* (Garavaglia *et al*, 2012). The operon encoding transcription factor and genes associated with curli assembly and transport is strongly affected by inactivation of UMP biosynthetic genes, indicating a significant interplay between nucleotide metabolism and biofilm formation (Garavaglia *et al*, 2012). Notably, the inactivation of PyrE impairs curli production in *E. coli*, a phenomenon monitored through the binding between Congo-red dye and curli (Hammar *et al*, 1995). In our experiment, treatment of lumichrome resulted in the inhibitory effect on Congo-red binding in a concentration-dependent manner (**Figure 4j**). This observation is consistent with the reduced bacterial motility and biofilm formation in Δ*pyrE* mutant strain of *Pseudomonas aeruginosa* (PA01) (Niazy *et al*, 2022) and *E. coli* (Lozano-Terol *et al*, 2023). Additionally, inhibition of biofilm formation in *E. coli* was observed with a decrease in PyrE activity due to acetylation at lysine 26 and 103 position (Lozano-Terol *et al*, 2023).

While the exact physiological significance requires further investigation, our findings indicate that the inhibitory effect of lumichrome on PyrE could contribute to the regulation of biofilm formation in bacteria. In this vein, previous studies have reported two noteworthy observations. Firstly, riboflavin and lumichrome have been identified as signaling molecules in bacterial quorum sensing (QS) and plant-microbe interactions (Rajamani *et al*, 2008; Dakora *et al*, 2015). Secondly, it has been reported that blocking the pyrimidine biosynthetic pathway affects riboflavin production in the filamentous hemiascomycete *Ashbya gossypii* (Silva *et al*, 2019). Considering these findings, we hypothesize that lumichrome may function as a modulator for these interconnected pathways, including pyrimidine biosynthesis, riboflavin production, and QS, regulating the flux of metabolic intermediates. This regulation could be achieved through lumichrome’s interaction with PyrE, potentially explaining the inhibition of biofilm formation in bacteria, as QS serves as an upstream regulatory component in the biofilm production process. Furthermore, based on the co-elution of lumichrome and UMPS in the PROMIS datasets from Arabidopsis and yeast, we speculate that the regulatory interplay between riboflavin pyrimidine nucleotide metabolism may be evolutionarily conserved. Notably, plants, fungi, and many bacteria encode the complete enzymatic machinery for *de novo* riboflavin and pyrimidine biosynthesis (Averianova *et al*, 2020; Fischer *et al*, 2004), supporting our speculations.

### Conclusions and Future Prospects

Here, we present a PMI network for *E. coli* obtained from a CF-MS-based strategy integrating two orthogonal separations, SEC and IEX. The main advantage of CF-MS-based approaches to assessing PMIs is that they offer a direct and largely unbiased view of PMIs and retrieve interactions that likely would be missed by other methods, such as the lumichrome-PyrE pairing described here. Although lumichrome is a conserved metabolite in prokaryotic and eukaryotic organisms, its potential roles and mode of action are poorly understood. Lumichrome has been associated with quorum sensing in bacteria (Mattmann & Blackwell, 2010) and a symbiotic relationship between legume plants and *rhizobia* bacteria (Phillips *et al*, 1999). Lumichrome supplementation enhances plant growth in a concentration-dependent manner (Gouws *et al*, 2012). Considering that lumichrome cofractionates with UMPS in PROMIS experiments from Arabidopsis, assessing the conservation of lumichrome regulation and its role in plant growth will be interesting. Another example of a novel association uncovered by our PMI network is the interaction of leucine-containing dipeptides and the fatty acid metabolism enzyme, FabF. Potential roles of dipeptides regulating lipid metabolism are intriguing, and the FabF-Val-Leu binding data provide motivation for future analysis of the functional consequences of this interaction on FabF activity and fatty acid metabolism.

However, it is important to acknowledge that CF-MS is not without caveats. In addition to false positives resulting from coincidental co-elution, CF-MS is biased towards stable complexes and will likely miss transient interactions. It will also favor abundant proteins and metabolites accessible to MS identification and quantification. Integrating different chromatographies can assist in distinguishing between true and coincidental co-elution but can miss other interactions favored by one chromatographic condition. Hence, CF-MS derived PMI networks can capture only a subset of true interactions. As pointed out before, although we refer to the PROMIS and omniTICC-derived interactions as high-confidence compared to those derived from a single chromatography, all inferred PMIs are obviously putative and require independent validation. Because validation is time-consuming, prioritization is a critical step, as in any interactomics study. Prioritization may be based on prior knowledge or additional experiments. Here, we explored co-elution across multiple PROMIS datasets and the conserved nature of small-molecule binding events. A different, highly promising avenue is *in silico* modeling of the binding events (Thieme & Walther, 2022; Walther, 2023). We anticipate that binding predictions will become instrumental in refining CF-MS-derived PMI networks and selecting true binding events.

## Materials and Methods

### Bacterial Growth

*Escherichia coli* strain BW25113 (Baba *et al*, 2006) was grown on the M9 minimal medium. For each separation, 2 liters of culture were harvested at OD = 0.8 and then the cells were washed thrice with 50 mM ammonium bicarbonate and 1.5 mM MgCl_2_, pH 7.5 and flash frozen for future use.

### Size exclusion chromatography (SEC)

The bacterial pellet was suspended in the cold lysis buffer (5 mL per 0.5 L of the culture). Composition of the lysis buffer: 50 mM Ammonium Bicarbonate, pH 7.5; 1.5 mM MgCl_2_; 150 mM NaCl; 5 mM dithiothreitol (DTT); 1 mM phenylmethylsulfonyl fluoride (PMSF); 1× protease inhibitor cocktail (Sigma Aldrich, cat. no. P9599). Cells were lysed on ice with a sonication probe. The lysate was subjected to an initial centrifugation of 7000 × g, 4 °C for 10 min, and the acquired supernatant was centrifuged again with the same parameters. The supernatant was further ultracentrifuged for 1 h at 165052 × g, 4 °C. The supernatant was concentrated on a Amicon centrifugation filter (Millipore-Sigma) with the 10 kDa cutoff to about 2.5 mL. The supernatant was transferred to a new falcon tube and centrifuged again at 18000 × g, 4 °C for 15 min to remove protein aggregates. Protein concentration was estimated with a Bradford assay. The lysate was diluted to 20 mg/mL and 2 mL were taken for the SEC separation. SEC was performed with a SRT-10 SEC-300 (Sepax) column connected to a NGC Quest 10 Chromatography System (Bio Rad) operating at 4 °C. Composition of the SEC buffer: 50 mM Ammonium Bicarbonate, pH 7.5; 1.5 mM MgCl_2_; 150 mM NaCl). The flow rate was set to 7 mL/min. We collected a total of 48 fractions containing 40-80 mL of elution. The fractions were lyophilized and stored at −80 °C for proteomic and metabolomic analysis. Detailed protocol can be find in (Sokolowska *et al*, 2019)

### Ion exchange chromatography (IEX)

The bacterial pellet was suspended in the cold lysis buffer (5 ml per 0.5 L of the culture). Cells were lysed on ice with a sonication probe. Composition of the lysis buffer: 50 mM Ammonium Bicarbonate, pH 7.5; 1.5 mM MgCl_2_; 5 mM dithiothreitol; 1 mM phenylmethylsulfonyl fluoride; 1× protease inhibitor cocktail (Sigma Aldrich, cat. no. P9599). The lysate was subjected to an initial centrifugation at 7000 × g, 4 °C for 10 min, and the acquired supernatant was centrifuged again with the same parameters. The supernatant was further ultracentrifuged for 1 h at 165052 × g, 4 °C. The supernatant was concentrated on the Amicon centrifugation filter (Milllipore-Sigma) with the 10 kDa cutoff to about 5 ml, after which 15 ml lysis buffer was added and concentrated again. The supernatant was transferred to a new falcon tube and centrifuged again at 18000 × g, 4 °C for 15 min to remove protein aggregates. Protein concentration was estimated with a Bradford assay. The sample concentration was diluted to 10 mg/ml and 2 ml were taken for the IEX sample separation. For the control separation, the 2.5 ml of 10 mg/ml protein lysate was denatured in 95 °C for 15 min and subsequently centrifuged in 18000 × g, 4 °C for 15 min. The supernatant was filtered through the Amicon centrifugation filter (Milllipore-Sigma) with the 10 kDa cutoff to allow all of the sample through. The flow-through was collected for IEX control separation. 2.0 mL of soluble fraction corresponding to 20 mg of protein, or control metabolite only sample, were used for the separations. IEX was performed with a Enrich Q, 5/50 mm (Bio Rad) column connected to a NGC Quest 10 Chromatography System (Bio Rad) operating at 4 °C. Composition of the IEX buffers; Buffer A: 50 mM Ammonium Bicarbonate, pH 7.5; 1.5 mM MgCl_2_, Buffer B: 50 mM Ammonium Bicarbonate, pH 7.5, 1.5 mM MgCl_2_, 1 M NaCl. The flow rate was set to 0.9 ml/min. Column was equilibrated with Buffer A. Elution gradient was set from 0% to 100% buffer B over the 20 ml. The collected 1 ml fractions were lyophilized and stored at −80 °C for proteomic and metabolomic analysis.

### Metabolite and protein extraction

The extraction protocol was adapted and modified from (Veyel *et al*, 2018). Proteins and metabolites were extracted from the lyophilized fractions using a methyl tert-butyl ether (MTBE)/methanol/water solvent system. Equal volumes of the polar fraction and protein pellet were dried in a centrifugal evaporator and stored at − 80 °C until they were processed further.

### Proteomics

IEX, PROMIS Rep1: Sample preparation, measurements and data analysis was performed by Biogenity ApS, Aalborg, Denmark. 40 lyophilized SEC fractions and 34 lyophilized IEX fractions were resuspended in 50µL Lysis buffer (consisting of 6 M Guanidinium Hydrochloride, 10 mM TCEP, 40 mM CAA, 50 mM HEPES pH 8.5). The samples were boiled at 95 °C for 5 minutes, followed by 5 minutes of sonication. The protein concentration of the fractions was determined with Pierce Rapid Gold BCA (Thermo), and 10 ug were taken forward for digestion. Samples were diluted 1:3 with 10% Acetonitrile, 50 mM HEPES pH 8.5. LysC (MS grade, Wako) was added in a 1:50 (enzyme to protein) ratio, and samples were incubated at 37 °C for 4 h. Samples were further diluted to a final 1:10 with 10% Acetonitrile, 50 mM HEPES pH 8.5. Trypsin (MS grade, Sigma) was added in a 1:100 (enzyme to protein) ratio and samples were incubated overnight at 37°C. Enzyme activity was quenched by adding 2% trifluoroacetic acid (TFA) to a final concentration of 1%. The peptides were desalted on a SOLAµ SPE plate (Rappsilber *et al*, 2007) (HRP, Thermo). For each sample, the filters were activated with 200 ul of 100% Methanol (HPLC grade, Sigma), then 200ul of 80% Acetonitrile, 0.1% formic acid. The filters were subsequently equilibrated 2x with 200ul of 1% TFA, 3% Acetonitrile, after which the sample was loaded using centrifugation at 1,500x rpm. After washing the filters twice with 200 ul of 0.1% formic acid, the peptides were eluted into clean 1.5ml Eppendorf tubes using 40% Acetonitrile, 0.1% formic acid. The eluted peptides were concentrated in an Eppendorf Speedvac, and re-constituted in 20 ul of 2% Acetonitrile, 1% TFA containing iRT peptides (Biognosys). For the ion exchange fractions, 500 ng were loaded onto EvoSep stagetips according to the manufacturer’s protocol. For each IEX fraction, peptides were analyzed using the pre-set ’15 samples per day’ method on the EvoSep One instrument. Peptides were eluted over an 88-min gradient. For each SEC pool, peptides were loaded onto a 2cm C18 trap column (ThermoFisher 164705), connected in-line to a 15 cm C18 reverse-phase analytical column (Thermo EasySpray ES804A) using 100% Buffer A (0.1% Formic acid in water) at 750 bar, the Thermo EasyLC 1200 HPLC system, and the column oven operating at 30°C. Peptides were eluted over a 140 minute gradient going from 10 to 23 % of Buffer B (80% acetonitrile, 0.1% formic acid) in 95 minutes, then to 38% Buffer B in 30 minutes, then to 60% Buffer B in 5 minutes, ending with 10 minutes at 95% Buffer B. The flow was set to 250 nl/min. Spectra were acquired with an Orbitrap ExplorisTM 480 instrument (Thermo Fisher Scientific) FAIMS ProTM Interface (ThermoFisher Scientific) set to CV of -45 V. Full MS spectra were collected at a resolution of 120,000, with normalized AGC target of 300% or maximum injection time of 45 ms and a scan range of 345–1500 *m/z*. AGC target for the DIA experiment was set to 1000%. The precursor mass range was set to 350-1400 m/z with 48 isolation windows of 21.5 Da and 1 Da overlap. The resolution was set to 15,000 with maximum injection time set to auto and a normalized collision energy of 30%. MS performance was verified for consistency by running complex cell lysate quality control standard. Raw files from the DIA experiments were analyzed using Spectronaut 15.5TM (Biognosys) using the BGS Factory Settings. PROMIS Rep2: Protein pellets were processed as described in (Thirumalaikumar *et al*, 2023). Briefly, protein pellets were dissolved in a denaturation buffer (6 M urea, 2 M thiourea dissolved in 50 mM Ammonium bicarbonate). Bradford assay was used to calculate protein concentration. 50 μg protein was taken for protein digestion. The protein extracts were reduced by the addition of DTT (100 μM) for 60 min at room temperature. Alkylation step was done using (300 μM) iodoacetamide for 60 min at dark. Residual iodoacetamide was quenched by additional (100 μM) DTT for 10 min. Finally, proteins were digested using trypsin/Lys-C mixture (Mass Spec Grade, Promega) for 16 h according to the manufacturer’s instruction. Digested peptide samples were desalted using C18 sep-pak column plates as described in (Thirumalaikumar *et al*, 2023). The eluted peptides were transferred to Eppendorf low-bind tubes and then dried using speed vac. The dried peptides were resuspended using a resuspension buffer (5% acetonitrile in 0.1% formic acid). Approximately, 1μg of the peptides were injected for analysis. The peptide mixtures were separated using a nano-liquid-chromatography system (Dionex ultimate 3000) using an acclaim pepmap C_18_ column. A flow rate of 300 nL/min was used for the complex peptide separation. The solvent A/B gradient were as follows: being isocratic at 3% B for 5 min, linearly increasing to 25% B at 80 min, linearly increasing to 45% B at 105 min, keeping at 95% B from 120 min to 131 min, shifting back to 3% B in 0.1 min and holding until 131.1 min. The peptide samples were sprayed using a nano bore stainless-steel emitter (Fisher Scientific). Peptides were analyzed using an Orbitrap-Exploris-480 MS™ mass spectrometer. Data was collected using a data-dependent acquisition (DDA) mode using a cycle time of 3 seconds. Standard mass spectrometer parameters were kept as described in (Henneberg *et al*, 2023), briefly: positive ion voltage ∼2.3 kV, ion transfer tube temperature at 320 °C; full scan orbitrap resolution 60,000, scan range *m*/*z* 350-1650, RF lens at 40%, maximum injection time mode was kept at Auto, AGC target was kept at standard; ddMS^2^ filters include monoisotopic peak determination for peptide, intensity threshold was kept at minimum intensity of 5000, charge state 2-8, exclude isotopes; ddMS^2^ isolation window *m*/*z* 1.6, HCD normalized collision energies 30%, collision energy type was kept at normalized, orbitrap resolution 15,000, RF lens 50 %, standard AGC target, auto maximum injection time. HeLa digests (Pierce, 88329) have been used to monitor the retention time drift and mass accuracy of the LC before and after each experiment. Raw data were analyzed using the Proteome Discoverer (version 2.5, ThermoFisher scientific) following the manufacturer’s instruction. PD Search was made using an *E. coli* protein database downloaded from Uniport. Common contaminants were compiled and added to the search. SEQUEST HT was used to assign the peptides, allowing a maximum of 2 missed tryptic cleavages. In addition, precursor mass tolerance of 10 ppm and a fragment mass tolerance of 0.02 Da, with a minimum peptide length of 6 AAs were selected. Cysteine-carbamidomethylation and methionine-oxidation were selected in default modifications. Label-free quantification (LFQ) based on MS1 precursor ion intensity was performed in Proteome Discoverer with a minimum Quan value threshold set to 0.0001 for unique peptides. The ‘3 Top N’ peptides were used for area calculation.

### Metabolomics

After extraction, the dried aqueous metabolites were measured using ultra-performance liquid chromatography (UPLC) coupled with a Q-Exactive mass spectrometer (Thermo Fisher Scientific) in positive and negative ionization modes, as described earlier (Veyel *et al*, 2018). Expressionist Refiner MS 12.0 (Genedata AG, Basel, Switzerland) was used for processing the LC–MS data. In-house library of authentic reference compounds was used to identify molecular features allowing 10 ppm mass deviation and dynamic retention time deviation (maximum 0.1 min).

### Data Analysis

Elution profiles were analyzed using standard settings in the PROMIsed app (Schlossarek *et al*, 2021), including pre-processing, deconvolution, and integration steps. The omniTICC profiles are derived from the manual curation of the IEX profiles coming from the control and sample preparations. Specifically, elution maxima corresponding to the free metabolites, based on the control separation, were removed for downstream analysis. Note that, however, as differential elution was primarily dictated by shift in the elution maxima, we also accepted elution maxima with the average fold-change intensity in the sample at least 3-time more than in the control.

### Protein purification

Recombinant PyrE was purified from an *E. coli* expression strain received from ASKA collection (Kitagawa *et al*, 2005) engineered to express a His-tagged PyrE. The process began with a bacterial preculture in LB medium, which was incubated at 37 °C for 3 hours. Protein expression was then induced by adding 100 μM of IPTG, followed by incubation for 4-5 hours at 37 °C. Frozen bacterial cells were used for subsequent protein purification. For small-scale purification, we utilized the His-Bind purification kit (Merck, USA) followed by manufacturer’s protocol. For large-scale purification, the supernatant was loaded onto a 1 ml HisTrap (Cytiva) column connected to NGC Quest 10 (Bio-Rad), previously equilibrated with Buffer A (25 mM Tris–HCl pH 8.0, 300 mM NaCl, 5% (v/v) glycerol, 10 mM imidazole). The column was washed with 10 ml of Buffer A and the recombinant protein was eluted with a linear gradient of imidazole (10– 300 mM, 50 ml). The fractions containing the enzyme of interest were collected and concentrated and desalted using ultra-centrifugal filters. The purified protein was stored in 0.1 M Tris-HCl buffer pH 8.8 at 4 °C for use in future experiments. FabF cloned into pET-28a expression vector was expressed in *E coli* BL21 (DE3) strains. Bacteria were grown in the LB medium to the OD of 0.8 - 1.0. Protein expression was then induced by adding 100 μM of IPTG, followed by 4 h incubation at RT. Expression, induction and purification of the recombinant protein were performed as described above. The purified protein was stored at 4 °C in 50 mM sodium phosphate buffer pH 7.4 supplemented with 300 mM NaCl.

### Microscale thermophoresis (MST)

A dilution series of PRPP and lumichrome was prepared by using 16-step, two-fold serial dilution for each ligand. The concentration in this series started at 781 μM for PRPP and 200 μM for lumichrome, decreasing with each step. These dilutions were carried out in their respective binding buffers, which consisted of 50 mM Tris-HCl (pH 8.0 for PRPP and pH 9.5 for lumichrome), 150 mM NaCl, 10 mM MgCl_2_, and 0.05% Tween 20. Each reaction mixture was created by mixing 6 μL of a protein working solution, containing 6.25 μM of His-tagged PyrE labeled with RED dye (Nanotemper Technologies, USA), with an equal volume of the respective ligand solution. The mixtures were loaded into standard capillaries. Subsequently, these capillaries were inserted into the Monolith device (Nanotemper Technologies, USA), and the measurement of MST was conducted following the manufacturer’s provided instructions. Analysis of MST data involved standard MST traces and the plotting of changes in normalized fluorescence (*F_norm_*) against the ligand concentration, enabling the determination of the dissociation constant (*K_d_*). This analysis was performed using the Monolith analysis software. FabF protein was labeled with the RED-MALEIMIDE or RDE-NHS dye (Nanotemper Technologies, USA) according to the manufacturer’s instructions. Binding was performed in premium capillaries in 25m M PIPES pH 7.5 and 200 mM NaCl. MST power was set to high. Data analysis was performed using Monolith software.

### Thermal shift assay (TSA)

To assess the shift in the unfolding temperature of PyrE in the presence of the ligands compared to the unfolding temperature obtained in the absence of any ligand, we conducted a thermal shift assay (TSA). Each reaction solution was prepared with 2.5 μg of purified protein, the ligand solution, and 1X protein thermal shift dye (Applied Biosystem, USA), following the manufacturer’s instructions. The melting curve for each treatment was monitored using a real-time PCR instrument, as the temperature was incrementally raised from 25 °C to 99 °C at a rate of 0.05 °C per second.

### Orotate phosphoribosyltransferase (OPRTase) assay

The activity of orotate phosphoribosyltransferase (OPRTase) was measured as previously described with some modification (SHIMOSAKA et al., 1985). The reaction mixture consisted of 100 mM Tris-HCl (pH 8.8), 5 mM MgCl_2_, 0.2 mM orotic acid and PRPP ranging from 0.125 to 4 mM. To examine the effect of lumichrome on PyrE activity, 100 μM of lumichrome was added. The reaction is initiated by adding PyrE into the mixture. The decrease in absorbance at 295 nm, resulting from the conversion of orotic acid (ε_295_= 3950 M^-1^ cm^-1^), was monitored after 1-minute of incubation at 30 °C. The activity was determined by calculating the amount of enzyme required to convert 1 μmol of orotic acid to orotidine monophosphate (OMP) per minute.

### Congo-red binding assay

*E. coli* W3110S was cultured overnight at 37 °C in LB medium. Next, 2 μL of the culture was inoculated onto LB 1/4 agar medium containing 0.004% Congo-red and 0.002% Coomassie blue (CR medium). To investigate the impact of lumichrome on Congo-red binding,10 μM and 100 μM of lumichrome was added into CR medium. The bacterial strain was grown at 30 °C for 24 h, followed by incubation at 4 °C for an additional 24 h. The bacterial cells were harvested and extracted using DMSO. After centrifugation at 13,500 x *g* for 1 min, the absorbance of the supernatant was measured at 497 nm.

### Molecular docking

3D structural information files for the proteins of interest were downloaded from RCSB PDB (Berman *et al*, 2000). Potential ligand binding sites for these structures were predicted using PrankWeb (Jakubec *et al*, 2022). If more than one binding pocket was predicted with high probability, the binding pocket with highest probability was chosen for the subsequent molecular docking. The chosen binding sites were then used to perform molecular docking with the SeeSAR software v13.0.1 (BioSolveIT GmbH). Subsequently, 2D interaction diagrams were generated for the top docking pose for each compound-protein complex using PoseView (Fricker *et al*, 2004).

## Supplementary material

**Supplementary Dataset S1.** Protein elution profiles, complete dataset.

**Supplementary Dataset S2.** Metabolite elution profiles, complete dataset.

**Supplementary Dataset S3.** IEX control, sample and omniTICC profiles for the subset of annotated metabolites.

**Supplementary Dataset S4a.** Protein elution profiles, input for PROMISed.

**Supplementary Dataset S4b.** Metabolite elution profiles, input for PROMISed.

**Supplementary Dataset S5a.** PMI network delineated using standard PROMISed app setting PCC >0.8 in PROMIS and omniTICC.

**Supplementary Dataset S5b.** PMI network delineated using standard PROMISed app setting PCC >0.7 in PROMIS and omniTICC.

**Supplementary Figure 1. Elution profiles.** a) Elution profiles of co-factors NADH and FAD. b) Elution profiles of hypoxanthine and its known interactor, purR. The red line indicates co-elution.

## Figures and Figure Legends

**Supplementary Figure 1.**
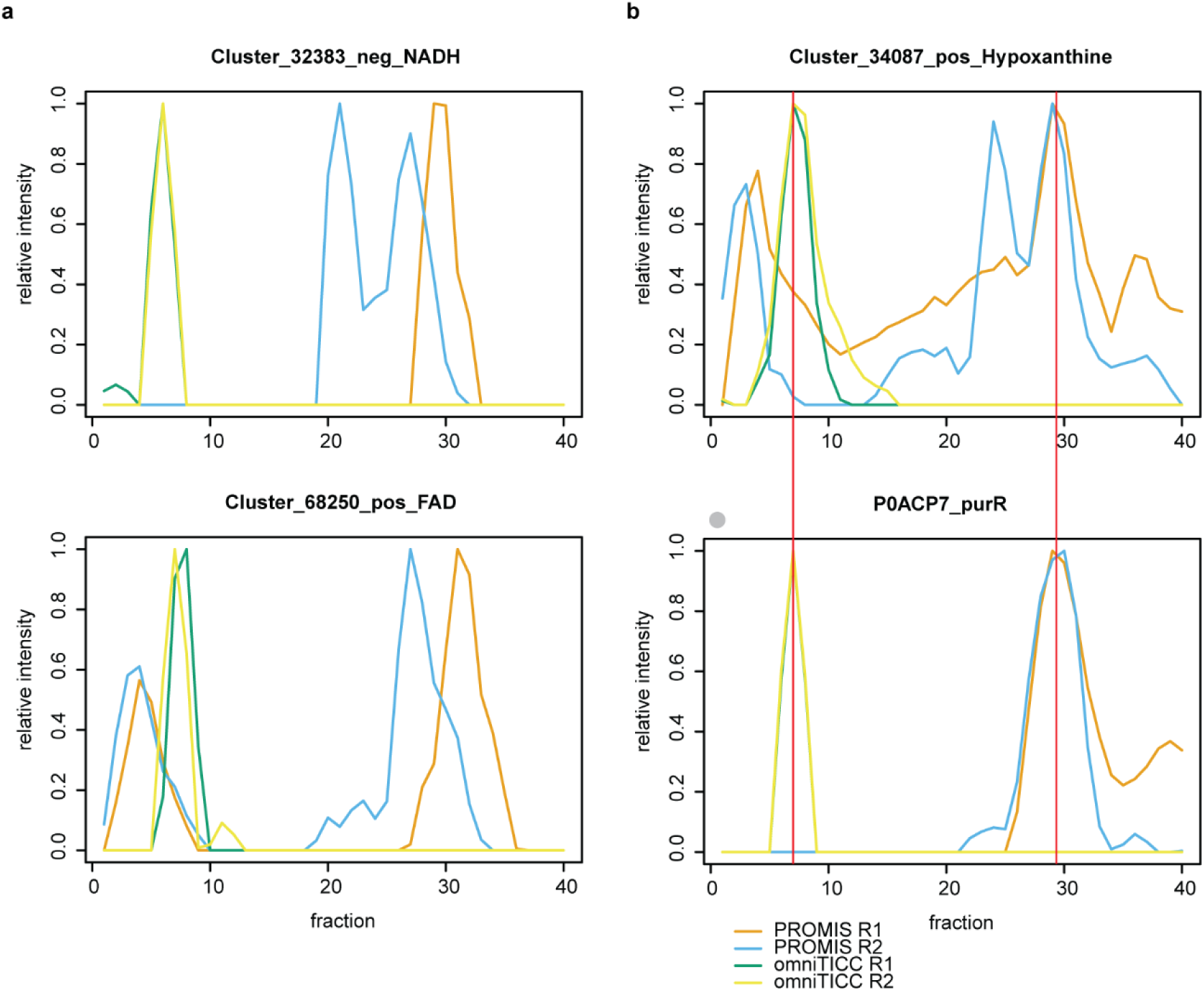
Elution profiles. a) Elution profiles of co-factors NADH and FAD. b) Elution profiles of hypoxanthine and its known interactor, purR. The red line indicates co-elution.

## Acknowledgements

We are grateful to Änne Michaelis for excellent technical assistance. The authors would like to acknowledge the support from Boyce Thompson Institute, Cornell University, Michigan State University and Max-Planck-Society. We are thankful to Biogenity DTU Proteomics Core for performing the proteomic sample measurements and contribution to data analysis. This work was supported by the U.S. National Science Foundation (GRANT 2226270 awarded to A.S.).

## Author Contributions

M.W. designed the experiments, conducted experiments. J.K. designed the experiments, conducted experiments, and wrote the manuscript. C.M. conducted experiments. V.T. performed proteomics measurements and analyzed proteomics data.

M.G. analyzed proteomics data. R.I.M. established the FPLC protein purification method. C.F.P. prepared proteomics samples. J.Z. and K.D. provided FabF protein expression construct and revised the manuscript. F.C.S. assisted in data analysis and writing the manuscript. H.Z. and D.W. performed docking analysis. A.S. designed experiments, analyzed data, wrote the manuscript, and supervised the study.

## Competing interest

F.C.S. is a founder of, consultant, and a stockholder for Ascribe Bioscience and Holoclara Inc. Other than that, the authors declare that they have no competing interests.

## Data availability

PROMISed app is available at https://github.com/DennisSchlossarek/PROMISed. Raw and analyzed proteomics and metabolomics data are available in a supplementary material. Raw chromatograms for proteomics were submitted to MassIVE repository (Perez-Riverol *et al*, 2022) (MSV000093709; MSV000093733; MSV000094003; MSV000094057). Raw chromatograms for metabolomics were submitted to Metabolights (Yurekten *et al*, 2023); study number MTBLS9098 (under curation).

## References

Aryal UK, Xiong Y, McBride Z, Kihara D, Xie J, Hall MC & Szymanski DB (2014) A proteomic strategy for global analysis of plant protein complexes. Plant Cell 26: 3867–3882

Averianova LA, Balabanova LA, Son OM, Podvolotskaya AB & Tekutyeva LA (2020) Production of Vitamin B2 (Riboflavin) by Microorganisms: An Overview. Front Bioeng Biotechnol 8: 570828

Baba T, Ara T, Hasegawa M, Takai Y, Okumura Y, Baba M, Datsenko KA, Tomita M, Wanner BL & Mori H (2006) Construction of Escherichia coli K-12 in-frame, single-gene knockout mutants: the Keio collection. Mol Syst Biol 2: 2006.0008

Baker SA & Rutter J (2023) Metabolites as signalling molecules. Nat Rev Mol Cell Biol 24: 355–374

Berman HM, Westbrook J, Feng Z, Gilliland G, Bhat TN, Weissig H, Shindyalov IN & Bourne PE (2000) The Protein Data Bank. Nucleic Acids Res 28: 235–242

Chan JNY, Vuckovic D, Sleno L, Olsen JB, Pogoutse O, Havugimana P, Hewel JA, Bajaj N, Wang Y, Musteata MF, et al (2012) Target identification by chromatographic co-elution: monitoring of drug-protein interactions without immobilization or chemical derivatization. Mol Cell Proteomics 11: M111.016642

Cheng L, Tanaka M, Yoshino A, Nagasato Y, Takata F, Dohgu S & Matsui T (2023) A memory-improving dipeptide, Tyr-Pro, can reach the mouse brain after oral administration. Sci Rep 13: 16908

Cho B-K, Federowicz SA, Embree M, Park Y-S, Kim D & Palsson BØ (2011) The PurR regulon in Escherichia coli K-12 MG1655. Nucleic Acids Res 39: 6456–6464

Chubukov V, Gerosa L, Kochanowski K & Sauer U (2014) Coordination of microbial metabolism. Nat Rev Microbiol 12: 327–340

Dakora FD, Matiru VN & Kanu AS (2015) Rhizosphere ecology of lumichrome and riboflavin, two bacterial signal molecules eliciting developmental changes in plants. Front Plant Sci 6: 700

Diether M, Nikolaev Y, Allain FH & Sauer U (2019) Systematic mapping of protein-metabolite interactions in central metabolism of Escherichia coli. Mol Syst Biol 15: e9008

Diether M & Sauer U (2017) Towards detecting regulatory protein–metabolite interactions. Curr Opin Microbiol 39: 16–23

Du F, Navarro-Garcia F, Xia Z, Tasaki T & Varshavsky A (2002) Pairs of dipeptides synergistically activate the binding of substrate by ubiquitin ligase through dissociation of its autoinhibitory domain. Proc Natl Acad Sci U S A 99: 14110– 14115

Edwards P, Nelsen JS, Metz JG & Dehesh K (1997) Cloning of the fabF gene in an expression vector and in vitro characterization of recombinant fabF and fabB encoded enzymes from Escherichia coli. FEBS Lett 402: 62–66

Fischer M, Römisch W, Saller S, Illarionov B, Richter G, Rohdich F, Eisenreich W & Bacher A (2004) Evolution of vitamin B2 biosynthesis: structural and functional similarity between pyrimidine deaminases of eubacterial and plant origin. J Biol Chem 279: 36299–36308

Fricker PC, Gastreich M & Rarey M (2004) Automated drawing of structural molecular formulas under constraints. J Chem Inf Comput Sci 44: 1065–1078

Garavaglia M, Rossi E & Landini P (2012) The pyrimidine nucleotide biosynthetic pathway modulates production of biofilm determinants in Escherichia coli. PLoS One 7: e31252

Ge SX, Jung D & Yao R (2020) ShinyGO: a graphical gene-set enrichment tool for animals and plants. Bioinformatics 36: 2628–2629

Gouws LM, Botes E, Wiese AJ, Trenkamp S, Torres-Jerez I, Tang Y, Hills PN, Usadel B, Lloyd JR, Fernie AR, et al (2012) The plant growth promoting substance, lumichrome, mimics starch, and ethylene-associated symbiotic responses in lotus and tomato roots. Front Plant Sci 3: 120

Gruber CH, Diether M & Sauer U (2021) Conservation of metabolic regulation by phosphorylation and non-covalent small-molecule interactions. Cell Syst

Hackett SR, Zanotelli VRT, Xu W, Goya J, Park JO, Perlman DH, Gibney PA, Botstein D, Storey JD & Rabinowitz JD (2016) Systems-level analysis of mechanisms regulating yeast metabolic flux. Science 354

Hammar M, Arnqvist A, Bian Z, Olsén A & Normark S (1995) Expression of two csg operons is required for production of fibronectin-and congo red-binding curli polymers in Escherichia coli K-12. Mol Microbiol 18: 661–670

Havugimana PC, Hart GT, Nepusz T, Yang H, Turinsky AL, Li Z, Wang PI, Boutz DR, Fong V, Phanse S, et al (2012) A census of human soluble protein complexes. Cell 150: 1068–1081

Heidenreich E, Pfeffer T, Kracke T, Mechtel N, Nawroth P, Hoffmann GF, Schmitt CP, Hell R, Poschet G & Peters V (2021) A Novel UPLC-MS/MS Method Identifies Organ-Specific Dipeptide Profiles. Int J Mol Sci 22

Henneberg LT, Singh J, Duda DM, Baek K, Yanishevski D, Murray PJ, Mann M, Sidhu SS & Schulman B (2023) Activity-based profiling of cullin-RING ligase networks by conformation-specific probes. bioRxiv

Hu Y, Flockhart I, Vinayagam A, Bergwitz C, Berger B, Perrimon N & Mohr SE (2011) An integrative approach to ortholog prediction for disease-focused and other functional studies. BMC Bioinformatics 12: 357

Ichinose T, Moriyasu K, Nakahata A, Tanaka M, Matsui T & Furuya S (2015) Orally administrated dipeptide Ser-Tyr efficiently stimulates noradrenergic turnover in the mouse brain. Biosci Biotechnol Biochem 79: 1542–1547

Jakubec D, Skoda P, Krivak R, Novotny M & Hoksza D (2022) PrankWeb 3: accelerated ligand-binding site predictions for experimental and modelled protein structures. Nucleic Acids Res 50: W593–W597

Jerabek-Willemsen M, Wienken CJ, Braun D, Baaske P & Duhr S (2011) Molecular interaction studies using microscale thermophoresis. Assay Drug Dev Technol 9: 342–353

Kim M-K, Song H-E, Kim YS, Rho S-H, Im YJ, Lee JH, Kang GB & Eom SH (2003) Crystallization and preliminary X-ray crystallographic analysis of orotate phosphoribosyltransferase from Helicobacter pylori. Mol Cells 15: 361–363

Kitagawa M, Ara T, Arifuzzaman M, Ioka-Nakamichi T, Inamoto E, Toyonaga H & Mori H (2005) Complete set of ORF clones of Escherichia coli ASKA library (a complete set of E. coli K-12 ORF archive): unique resources for biological research. DNA Res 12: 291–299

Kosmacz M, Sokołowska EM, Bouzaa S & Skirycz A (2020) Towards a functional understanding of the plant metabolome. Current Opinion in Plant Biology 55: 47–51 doi:10.1016/j.pbi.2020.02.005 [PREPRINT]

Kuhn M, Szklarczyk D, Franceschini A, Campillos M, von Mering C, Jensen LJ, Beyer A & Bork P (2010) STITCH 2: an interaction network database for small molecules and proteins. Nucleic Acids Res 38: D552–6

Ledezma Tejeida DE, Schastnaya E & Sauer U (2021) Metabolism as a signal generator in bacteria.

Lee Y, Okita TW & Szymanski DB (2021) A co-fractionation mass spectrometry-based prediction of protein complex assemblies in the developing rice aleurone-subaleurone. Plant Cell 33: 2965–2980

Lee Y & Szymanski DB (2021) Multimerization variants as potential drivers of neofunctionalization. Sci Adv 7

Lempp M, Farke N, Kuntz M, Freibert SA, Lill R & Link H (2019) Systematic identification of metabolites controlling gene expression in E. coli. Nat Commun 10: 4463

Lim YT, Prabhu N, Dai L, Go KD, Chen D, Sreekumar L, Egeblad L, Eriksson S, Chen L, Veerappan S, et al (2018) An efficient proteome-wide strategy for discovery and characterization of cellular nucleotide-protein interactions. PLoS One 13: e0208273

Link H, Kochanowski K & Sauer U (2013) Systematic identification of allosteric protein-metabolite interactions that control enzyme activity in vivo. Nat Biotechnol 31: 357–361

Li Y, Kuhn M, Zukowska-Kasprzyk J, Hennrich ML, Kastritis PL, O’Reilly FJ, Phapale P, Beck M, Gavin A-C & Bork P (2021) Coupling proteomics and metabolomics for the unsupervised identification of protein–metabolite interactions in Chaetomium thermophilum. PLoS One 16: e0254429

Lozano-Terol G, Gallego-Jara J, Sola-Martínez RA, Ortega Á, Martínez Vivancos A, Cánovas Díaz M & de Diego Puente T (2023) Regulation of the pyrimidine biosynthetic pathway by lysine acetylation of E. coli OPRTase. FEBS J 290: 442– 464

Luzarowski M & Skirycz A (2019) Emerging strategies for the identification of protein– metabolite interactions. J Exp Bot 70: 4605–4618

Luzarowski M, Vicente R, Kiselev A, Wagner M, Schlossarek D, Erban A, de Souza LP, Childs D, Wojciechowska I, Luzarowska U, et al (2021) Global mapping of protein– metabolite interactions in Saccharomyces cerevisiae reveals that Ser-Leu dipeptide regulates phosphoglycerate kinase activity. Communications Biology 4: 181

Mallam AL, Sae-Lee W, Schaub JM, Tu F, Battenhouse A, Jang YJ, Kim J, Wallingford JB, Finkelstein IJ, Marcotte EM, et al (2019) Systematic Discovery of Endogenous Human Ribonucleoprotein Complexes. Cell Rep 29: 1351–1368.e5

Mattmann ME & Blackwell HE (2010) Small molecules that modulate quorum sensing and control virulence in Pseudomonas aeruginosa. J Org Chem 75: 6737–6746

McWhite CD, Papoulas O, Drew K, Cox RM, June V, Dong OX, Kwon T, Wan C, Salmi ML, Roux SJ, et al (2020) A Pan-plant Protein Complex Map Reveals Deep Conservation and Novel Assemblies. Cell 181: 460–474.e14

Minen RI, Thirumalaikumar VP & Skirycz A (2023) Proteinogenic dipeptides, an emerging class of small-molecule regulators. Curr Opin Plant Biol 75: 102395

Mizushige T, Uchida T & Ohinata K (2020) Dipeptide tyrosyl-leucine exhibits antidepressant-like activity in mice. Sci Rep 10: 2257

Moreno JC, Rojas BE, Vicente R, Gorka M, Matz T, Chodasiewicz M, Peralta-Ariza JS, Zhang Y, Alseekh S, Childs D, et al (2021) Tyr-Asp inhibition of glyceraldehyde 3-phosphate dehydrogenase affects plant redox metabolism. EMBO J 40: e106800

Naka K, Jomen Y, Ishihara K, Kim J, Ishimoto T, Bae E-J, Mohney RP, Stirdivant SM, Oshima H, Oshima M, et al (2015) Dipeptide species regulate p38MAPK-Smad3 signalling to maintain chronic myelogenous leukaemia stem cells. Nat Commun 6: 8039

Niazy AA, Lambarte RNA & Alghamdi HS (2022) de novo pyrimidine synthesis pathway inhibition reduces motility virulence of Pseudomonas aeruginosa despite complementation. J King Saud Univ Sci 34: 102040

Orsak T, Smith TL, Eckert D, Lindsley JE, Borges CR & Rutter J (2012) Revealing the Allosterome: Systematic Identification of Metabolite–Protein Interactions. Biochemistry 51: 225–232

Perez-Riverol Y, Bai J, Bandla C, García-Seisdedos D, Hewapathirana S, Kamatchinathan S, Kundu DJ, Prakash A, Frericks-Zipper A, Eisenacher M, et al (2022) The PRIDE database resources in 2022: a hub for mass spectrometry-based proteomics evidences. Nucleic Acids Res 50: D543–D552

Phillips DA, Joseph CM, Yang GP, Martinez-Romero E, Sanborn JR & Volpin H (1999) Identification of lumichrome as a sinorhizobium enhancer of alfalfa root respiration and shoot growth. Proc Natl Acad Sci U S A 96: 12275–12280

Piazza I, Kochanowski K, Cappelletti V, Fuhrer T, Noor E, Sauer U & Picotti P (2018) A Map of Protein-Metabolite Interactions Reveals Principles of Chemical Communication. Cell 172: 358–372.e23

Rajamani S, Bauer WD, Robinson JB, Farrow JM 3rd, Pesci EC, Teplitski M, Gao M, Sayre RT & Phillips DA (2008) The vitamin riboflavin and its derivative lumichrome activate the LasR bacterial quorum-sensing receptor. Mol Plant Microbe Interact 21: 1184–1192

Rappsilber J, Mann M & Ishihama Y (2007) Protocol for micro-purification, enrichment, pre-fractionation and storage of peptides for proteomics using StageTips. Nat Protoc 2: 1896–1906

Scapin G, Ozturk DH, Grubmeyer C & Sacchettini JC (1995) The crystal structure of the orotate phosphoribosyltransferase complexed with orotate and alpha-D-5-phosphoribosyl-1-pyrophosphate. Biochemistry 34: 10744–10754

Schäkermann S, Wüllner D, Yayci A, Emili A & Bandow JE (2021) Applicability of Chromatographic Co-Elution for Antibiotic Target Identification. Proteomics 21: e2000038

Schlossarek D, Luzarowski M, Sokołowska E, Górka M, Willmitzer L & Skirycz A (2021) PROMISed: A novel web-based tool to facilitate analysis and visualization of the molecular interaction networks from co-fractionation mass spectrometry (CF-MS) experiments. Comput Struct Biotechnol J 19: 5117–5125

Schlossarek D, Luzarowski M, Sokołowska EM, Thirumalaikumar VP, Dengler L, Willmitzer L, Ewald JC & Skirycz A (2022) Rewiring of the protein–protein–metabolite interactome during the diauxic shift in yeast. Cellular and Molecular Life Sciences 79 doi:10.1007/s00018-022-04569-8

Schlossarek D, Zhang Y, Sokolowska EM, Fernie AR, Luzarowski M & Skirycz A (2023) Don’t let go: co-fractionation mass spectrometry for untargeted mapping of protein-metabolite interactomes. Plant J 113: 904–914

Silva R, Aguiar TQ, Oliveira C & Domingues L (2019) Physiological characterization of a pyrimidine auxotroph exposes link between uracil phosphoribosyltransferase regulation and riboflavin production in Ashbya gossypii. N Biotechnol 50: 1–8

Skinnider MA, Scott NE, Prudova A, Kerr CH, Stoynov N, Stacey RG, Chan QWT, Rattray D, Gsponer J & Foster LJ (2021) An atlas of protein-protein interactions across mouse tissues. Cell 184: 4073–4089.e17

Sokolowska EM, Schlossarek D, Luzarowski M & Skirycz A (2019) PROMIS: Global analysis of PROtein-metabolite interactions. Curr Protoc Plant Biol 4: e20101

Tegeder M & Rentsch D (2010) Uptake and partitioning of amino acids and peptides. Mol Plant 3: 997–1011

Thieme S & Walther D (2022) Biclique extension as an effective approach to identify missing links in metabolic compound–protein interaction networks. Bioinformatics Advances 2

Thirumalaikumar VP, Fernie AR & Skirycz A (2023) Untargeted Proteomics and Metabolomics Analysis of Plant Organ Development. Methods Mol Biol 2698: 75–85

Thirumalaikumar VP, Wagner M, Balazadeh S & Skirycz A (2020) Autophagy is responsible for the accumulation of proteogenic dipeptides in response to heat stress in Arabidopsis thaliana. FEBS J

Tian M, von Dahl CC, Liu P-P, Friso G, van Wijk KJ & Klessig DF (2012) The combined use of photoaffinity labeling and surface plasmon resonance-based technology identifies multiple salicylic acid-binding proteins. Plant J 72: 1027–1038

Treadwell GE & Metzler DE (1972) Photoconversion of riboflavin to lumichrome in plant tissues. Plant Physiol 49: 991–993

Tsukahara T, Yamagishi S, Neyama H & Ueda H (2018) Tyrosyl-tRNA synthetase: A potential kyotorphin synthetase in mammals. Peptides 101: 60–68 doi:10.1016/j.peptides.2017.12.026

Venegas-Molina J, Molina-Hidalgo FJ, Clicque E & Goossens A (2021) Why and How to Dig into Plant Metabolite-Protein Interactions. Trends Plant Sci 26: 472–483

Veyel D, Kierszniowska S, Kosmacz M, Sokolowska EM, Michaelis A, Luzarowski M, Szlachetko J, Willmitzer L & Skirycz A (2017) System-wide detection of protein-small molecule complexes suggests extensive metabolite regulation in plants. Sci Rep 7: 42387

Veyel D, Sokolowska EM, Moreno JC, Kierszniowska S, Cichon J, Wojciechowska I, Luzarowski M, Kosmacz M, Szlachetko J, Gorka M, et al (2018) PROMIS, global analysis of PROtein–metabolite interactions using size separation in Arabidopsis thaliana. J Biol Chem 293: 12440–12453

Wagner M, Zhang B, Tauffenberger A, Schroeder FC & Skirycz A (2021) Experimental methods for dissecting the terra-incognita of protein-metabolite interactomes. Current Opinion in Systems Biology: 100403

Walther D (2023) Specifics of Metabolite-Protein Interactions and Their Computational Analysis and Prediction. Methods Mol Biol 2554: 179–197

Wan C, Borgeson B, Phanse S, Tu F, Drew K, Clark G, Xiong X, Kagan O, Kwan J, Bezginov A, et al (2015) Panorama of ancient metazoan macromolecular complexes. Nature 525: 339–344

Xia Z, Turner GC, Hwang C-S, Byrd C & Varshavsky A (2008) Amino acids induce peptide uptake via accelerated degradation of CUP9, the transcriptional repressor of the PTR2 peptide transporter. J Biol Chem 283: 28958–28968

Yurekten O, Payne T, Tejera N, Amaladoss FX, Martin C, Williams M & O’Donovan C (2023) MetaboLights: open data repository for metabolomics. Nucleic Acids Res

Zhang Z, Zhao Y, Wang X, Lin R, Zhang Y, Ma H, Guo Y, Xu L & Zhao B (2016) The novel dipeptide Tyr-Ala (TA) significantly enhances the lifespan and healthspan of Caenorhabditis elegans. Food Funct 7: 1975–1984

